# Competitive plasticity to reduce the energetic costs of learning

**DOI:** 10.1101/2023.04.04.535544

**Authors:** Mark C.W. van Rossum

## Abstract

The brain is not only constrained by energy needed to fuel computation, but it is also constrained by energy needed to form memories. Experiments have shown that learning simple conditioning tasks already carries a significant metabolic cost. Yet, learning a task like MNIST to 95% accuracy appears to require at least 10^8^ synaptic updates. Therefore the brain has likely evolved to be able to learn using as little energy as possible. We explored the energy required for learning in feedforward neural networks. Based on a parsimonious energy model, we propose two plasticity restricting algorithms that save energy: 1) only modify synapses with large updates, and 2) restrict plasticity to subsets of synapses that form a path through the network. Combining these two methods leads to substantial energy savings while only incurring a small increase in learning time. In biology networks are often much larger than the task requires. In particular in that case, large savings can be achieved. Thus competitively restricting plasticity helps to save metabolic energy associated to synaptic plasticity. The results might lead to a better understanding of biological plasticity and a better match between artificial and biological learning. Moreover, the algorithms might also benefit hardware because in electronics memory storage is energetically costly as well.

---

Energy availability is a vital necessity for biological organisms. The nervous system is a particularly intensive energy consumer. Though it constitutes approximately 2%of human body mass, it is responsible for roughly 20% of basal metabolism – continuously consuming some 20W (Attwell and Laughlin, 2001; Harris et al., 2012). Energy requirements of neural signaling are now widely seen as an important design constraint of the brain. Sparse codes and sparse connectivity (pruning) can be used to lower energy requirements (Levy and Baxter, 1996). Learning rules have been designed that yield energy efficient networks (Sacramento et al., 2015; Grytskyy and Jolivet, 2021).

However, learning itself is also energetically costly: in classical conditioning experiments with flies the formation of long-term memory reduced lifespan by 20% when the flies were subsequently starved (Mery and Kawecki, 2005). Moreover, starving fruit flies halt long-term memory formation; nevertheless forcing memory expression reduced their lifespan (Plaçais and Preat, 2013). We have estimated the cost of learning in fruitflies at 10mJ/bit (Girard et al., 2023). In mammals there is physiological evidence that energy availability gates long lasting forms of plasticity (Potter et al., 2010). Furthermore, in humans there is behavioral evidence for a correlation between metabolism and memory (Smith et al., 2011; Klug et al., 2022).

Given this evidence, we hypothesize that biological neural plasticity is designed to be energy efficient and that including energy constraints could lead to computational learning rules more closely resembling biology, and a better understanding of observed synaptic plasticity rules. As precise metabolic cost models are not yet available, we aim at algorithmic principles rather than biophysical mechanisms that increase energy efficiency. Specifically we examine how energy needed for plasticity can be reduced while overall learning performance can be maintained.

Our study is inspired by experimental observations that despite common fluctuations in synaptic strength, the number of synapses undergoing permanent modification appears restricted. First plasticity is spatially limited, e.g. during motor learning plasticity is restricted to certain dendritic branches (Cichon and Gan, 2015). Second, plasticity is temporally limited. E.g. it is not possible to induce late-phase LTP twice in rapid succession (Rioult-Pedotti et al., 2000; Kramár et al., 2012). This stands in stark contrast with traditional backprop which updates *every* synapse on *every* trial which, as we shall see, can lead to very inefficient learning.

We use artificial neural networks as model of neural learning. While neural networks trained with back-propagation are an abstraction of biological learning, it allows for an effective way to teach networks complex associations that is currently not matched by more biological networks or algorithms. Biological implementations of back-propagation have been suggested (e.g. Sacramento et al., 2018), but it is likely that these are less energy efficient as learning times are typically longer.

In artificial neural network research there is a similar interest in limiting plasticity, but typically for other reasons. Carefully selecting which synapses to modify can prevent overwriting previously stored information, also known as catastrophic forgetting (Sezener et al., 2021). Randomly switching plasticity on and off can help regularization, as in drop-out and its variants (Salvetti et al., 1994). Communication with the memory systems is also energetically costly in computer hardware. Strikingly, storing two 32bit numbers in DRAM is far more expensive than multiplying them (Han et al., 2016). Recent studies have started to explore algorithms that reduce these costs by limiting the updates to weight matrices (Han et al., 2016; Golub et al., 2018).

## Methods

### Data-set

As data set we use the standard MNIST digit classification task. Similar results were found on the fashionMNIST data set. The data were offset so that the mean input was zero. This common preprocessing step is important. Namely, when trained with backprop, the weight update to weight *w_ij_* between input *x_j_* to a unit with error *δ_i_* is

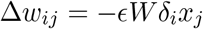

As the weight update is proportional to the input value *x_j_*, there is no plasticity for zero-valued inputs, even when *δ_i_* ≠ 0. However, this could be a confounding factor as we would like to be in full control of the number of plasticity events. After zero-meaning the inputs, only a negligible number of inputs will be exactly zero and this issue does not arise. Yet, the relative amount of achievable savings is not substantially affected by this assumption, Fig. 6.

**Figure 1:**
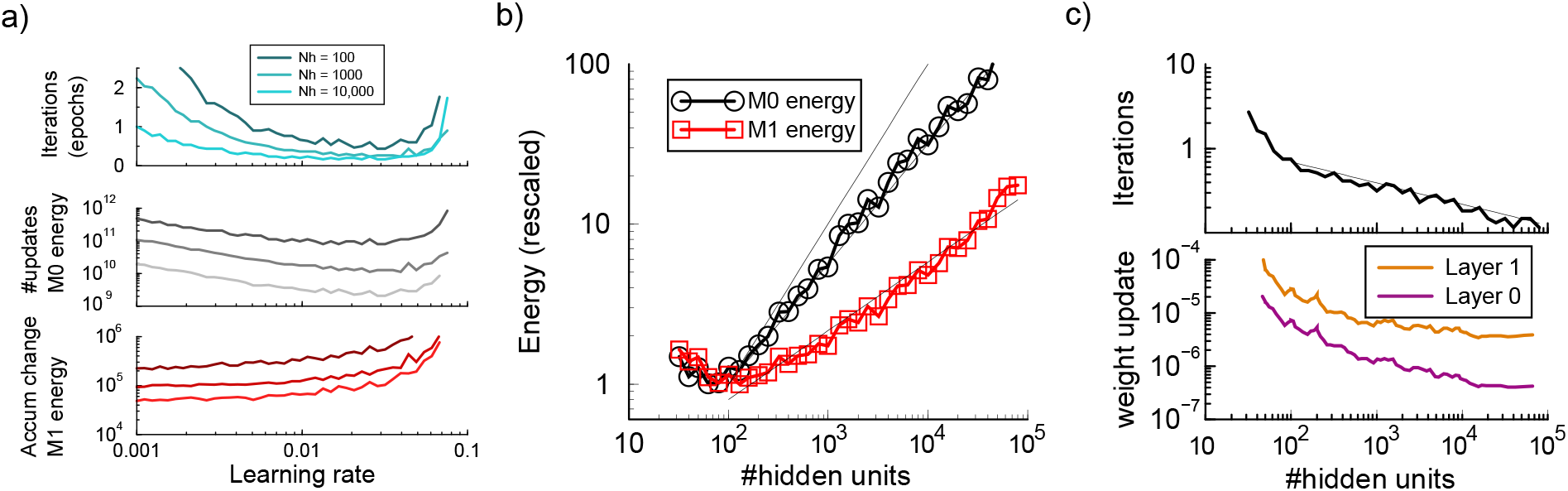
Energy requirements to train large standard backpropagation networks. a). Energy required to train a network with 100, 1000, or 10000 hidden units as a function of the learning rate. Top panel: number of iterations to reach 95% accuracy. Middle panel: The total number of updates (*M*_0_ energy) is minimal when the number of iterations is minimal. Bottom panel: The accumulated changes (*M*_1_ energy) is lowest in the limit of small learning rates. b). Energy measures versus network size. Since the network has one hidden layer, network size is expressed in the number of hidden units. Black curve is the number of updates (*M*_0_), red is the sum of all absolute changes (*M*_1_); both grow in large networks. The y-axis was scaled so that the minimum energy was one. The three thin lines represent slope one and powerlaw fits (see text). c). Learning speed and update size vs network size. Top: Large networks train somewhat faster to the 95% accuracy criterion than small networks. Bottom: the mean absolute weight update is smaller in large networks.

**Figure 2:**
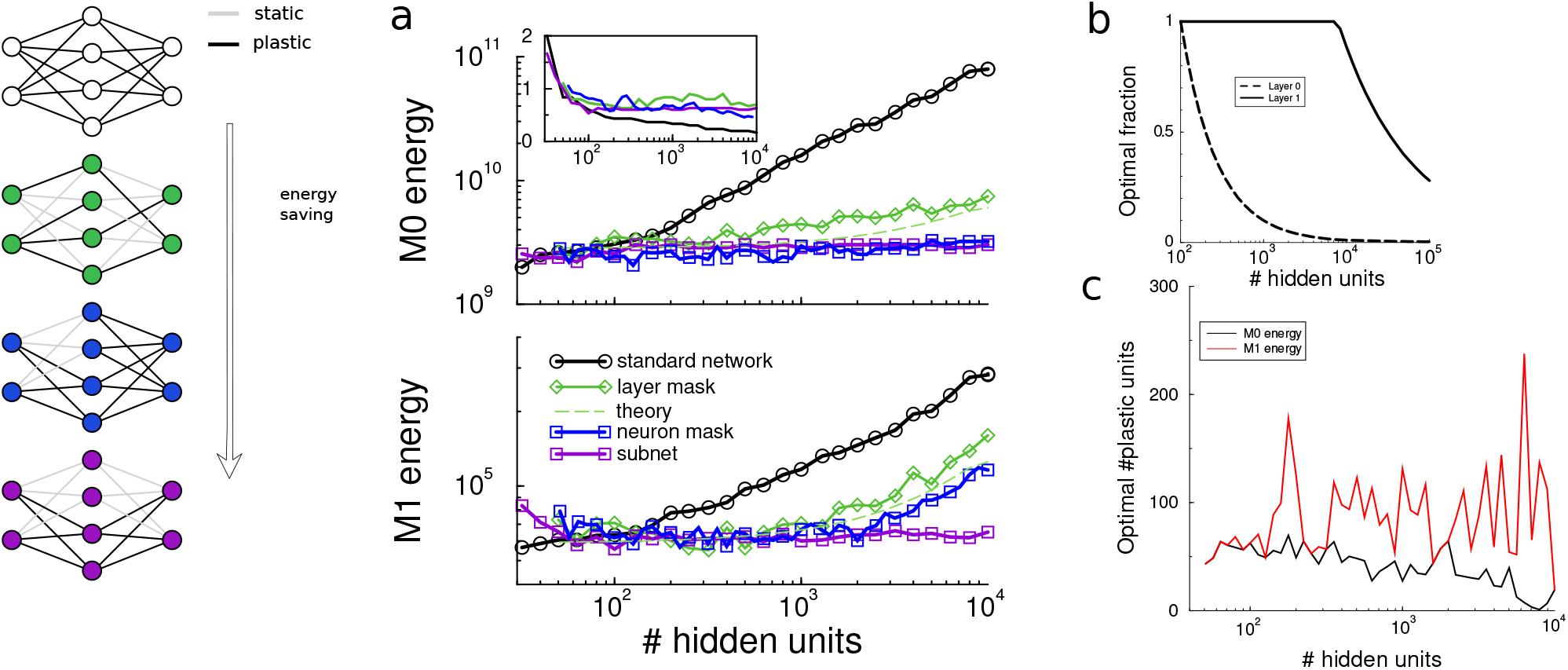
Restricting plasticity using random masks saves energy. a) M0 energy (total number of updates) and M1 energy (accumulated changes) for a standard network against network size (black). A random synaptic mask saves energy but cannot prevent an eventual increase in large networks (green; theory in dashed green). A bit more energy is saved when only a random subset of neurons in the hidden layer is plastic (blue). Most energy is saved when input and output plasticity is coordinated so that neurons with plastic inputs have plastic outputs, energy becomes independent of network size (purple). a-right) Diagram of the various masking settings. b) Theoretically optimal number of plastic synapses (used for green curve in panel a). c) The optimal number of plastic neurons found when using a neuron mask (corresponding to blue curve in a).

**Figure 3:**
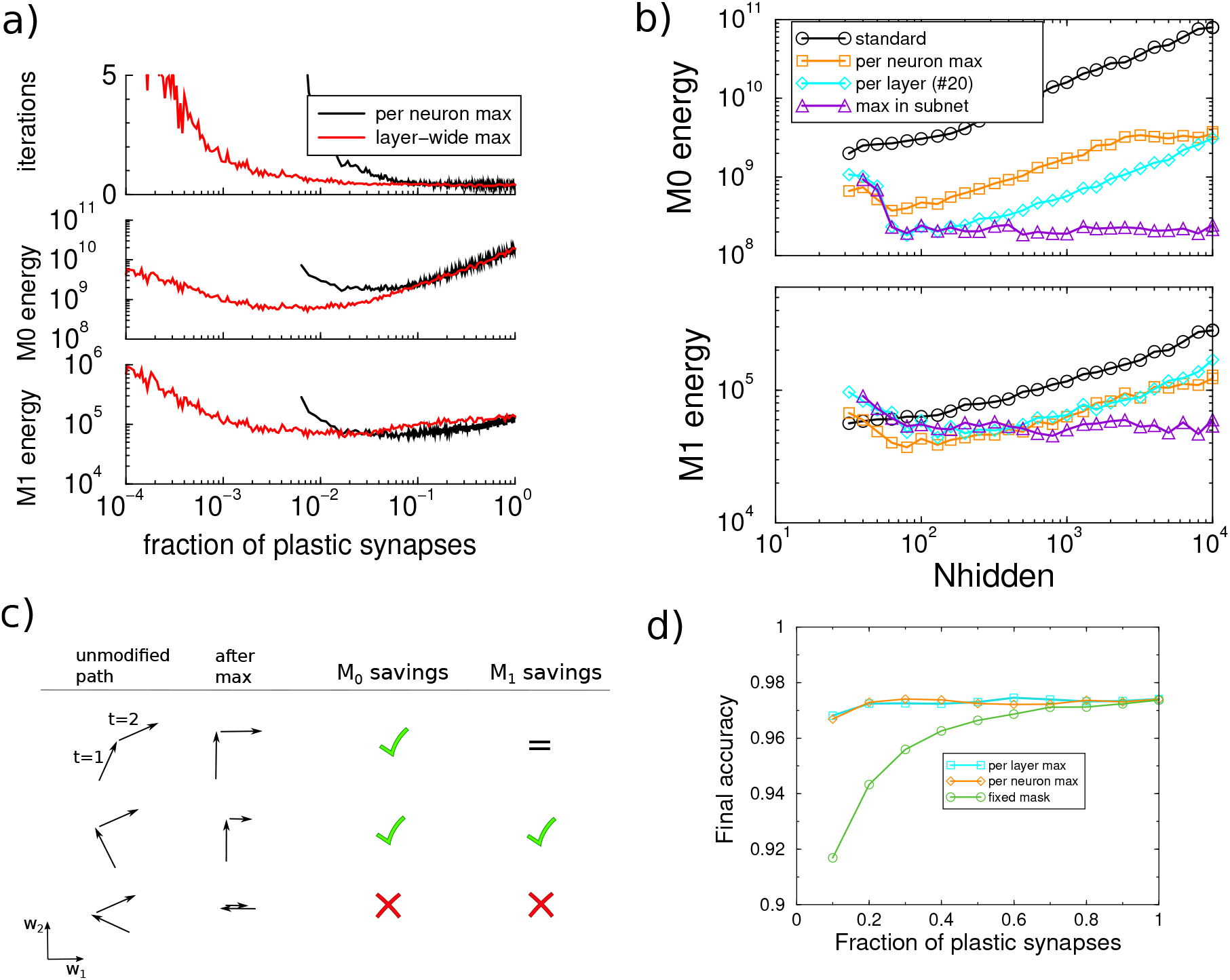
Energy required for network training when only synapses with large updates are modified. a) Iterations and energy in a network with 1000 hidden units as a function of the fraction of plastic synapses. The largest updates were selected across the layer (red) or for each hidden layer unit (black). b) Energy requirements as a function of the network size. Allowing only the largest synaptic updates per layer saves substantial amounts of energy (cyan curve), as did only allowing the largest updates per neuron (orange curve), compared to a standard network where all synapses are updated (black curve). The fraction of updated synapses was optimized for each network size. Further savings can be achieved by restricting the selection to subnets (purple), leading to a size independent energy need. c) Diagram of two subsequent weight updates in three hypothetical case (top to bottom). After the maximum selection, the updates are along the cardinal axes. In a particular when the weight trajectory does not return on itself (top), the selection saves *M*_0_ energy but *M*_1_ energy is identical. d) Test accuracy of a small network (100 hidden units) trained to saturation. A fixed mask limits performance, but the maximum selection does not.

**Figure 4:**
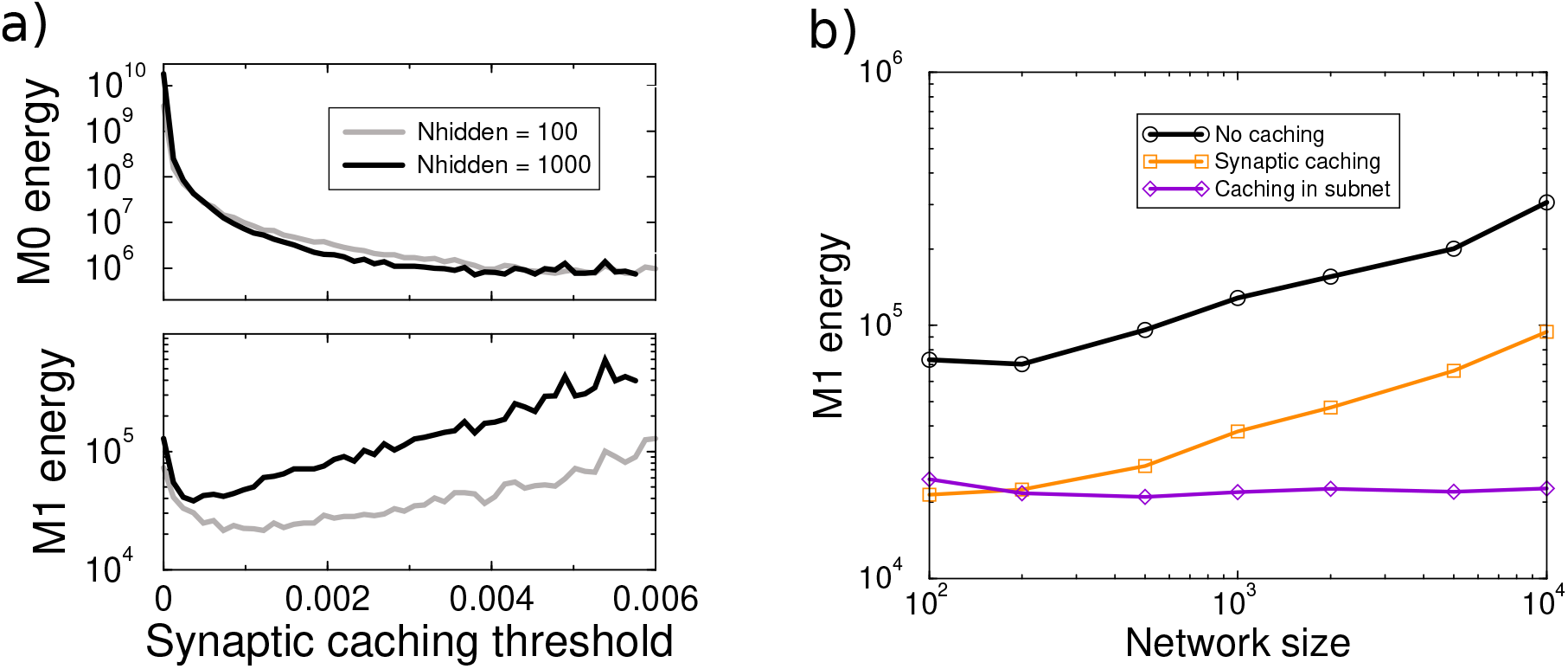
Synaptic caching. a) Energy savings achieved by synaptic caching for networks with 100 and 1000 hidden units. The *M*_0_ energy that only counts consolidation events, is minimal when the threshold is large, and is independent of network size. The *M*_1_ energy trades off cost transient plasticity with consolidation costs and learning time. b) Synaptic caching saves *M*_1_ energy. The *M*_1_ energy is made independent of size by restricting the plasticity to subnets.

**Figure 5:**
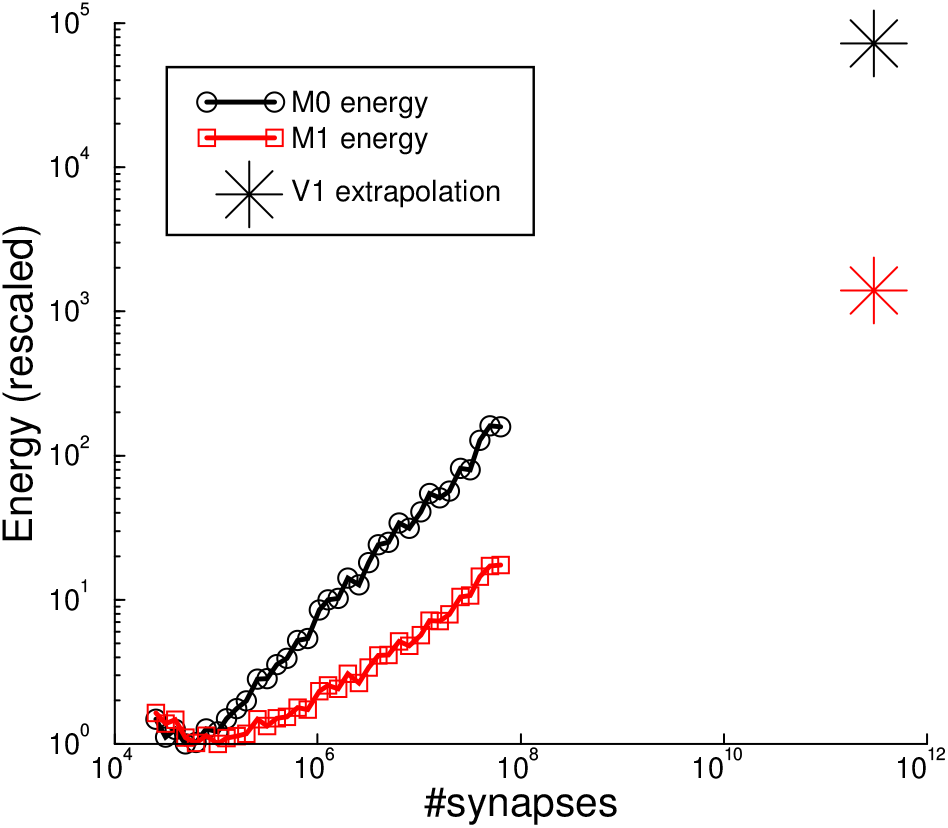
Extrapolation of results of Fig.1b to the number of synapses in macaque V1. Naive, unrestricted backprop learning would use 10^5^ × more synaptic updates and 10^3^ × more *M*_1_ energy than minimally required.

**Figure 6:**
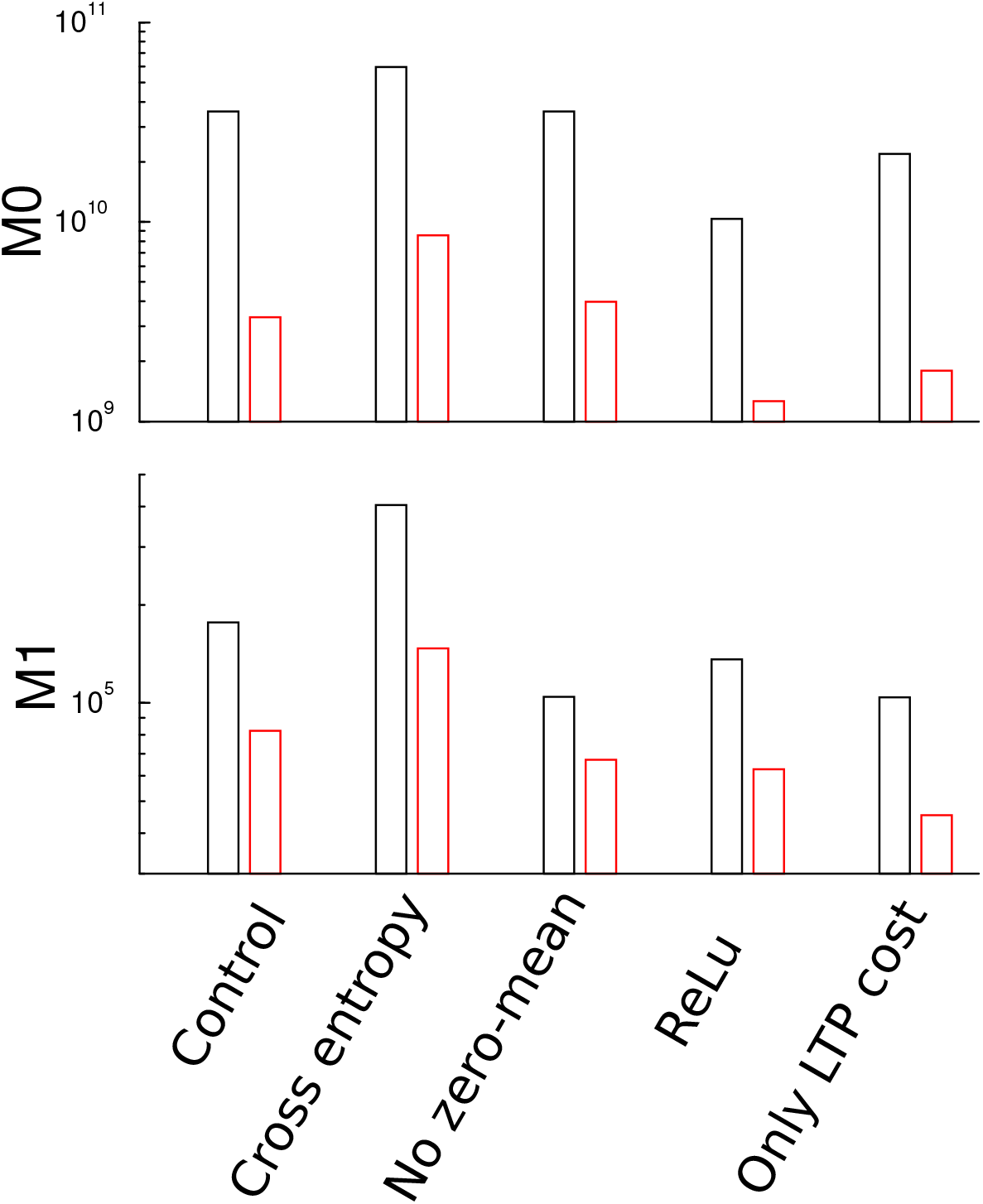
Influence of various assumptions on energy. Each pair of bars shows the energy required for a network with 2500 hidden units with unconstrained plasticity (black), and when using the optimal fraction of plastic synapses with a fixed mask (red). From left to right, **Control:** as in main text for comparison; **Cross entropy:** training on cross entropy loss function; **No zero-mean:** without zero-meaning the data; **ReLU:** using linear rectifying (ReLU) units in the hidden layer; **Only LTP cost:** only positive weight changes cost energy, negative changes are free. While the energy levels change, the amount of savings achievable remains similar.

### Network architecture

We used a network with 784 input units (equal to the number of MNIST pixels) with bias, a variable number of (non-convolutional) hidden units in a single hidden unit layer, and 10 output units (one for each digit). Networks had all-to-all, non-convolutional connectivity. The hidden layer units used a leaky rectifying linear activation function (lReLU), so that *g*(*x*) = *x* if *x* ≥ 0 and *g*(*x*) = *βx* with *β* = 0.1. A rectified linear activation function *g*(*x*) = max(0, *x*) would lead to a substantial fraction of neurons with zero activation in the hidden layer on a given sample and turn off plasticity of many synapses between hidden and output layer. As above, this would potentially be confounding, but it is avoided by using ‘leaky’ units. This did not substantially change the achievable savings, Fig. 6.

### Training

The activation of the ten output units is used to train the network. The target distribution was one-hot encoding of the image labels. For classification tasks it is common to apply a soft-max non-linearity *y_i_* = exp(*h_i_*)/ ∑*_j_* exp *h_j_*, where *h_i_* is the net input to each unit, so that the output activities represent a normalized probability distribution. One then trains the network by minimizing the cross entropy loss between the output distribution and the target distribution.

However, it is easy to see that when the target is a deterministic one-hot (0 and 1) distribution, the cross entropy loss only vanishes in the limit of diverging *h_i_*, which in turn requires diverging weights. As cross-entropy loss minimization is energy inefficient, we used linear output units and minimized the mean squared error between output and target. This did not substantially change the achievable savings, Fig. 6.

We used the backpropagation learning algorithm to train the network. Weights were initialized with small Gaussian values (*σ* = 0.01). In preliminary simulations, regularization was seen to have no significant effect on energy or savings, and was subsequently omitted. Samples were presented in a random order. Because batching would require storing synaptic updates across samples, which would be biologically challenging, the training was not batched and the synaptic weights were updated after every sample. We did not find substantial energy savings when using adaptive learning schemes such as ADAM. As such schemes would furthermore be hard to implement biologically, we instead fixed the learning rate.

Every 1000 samples training was interrupted and performance on a validation data set was measured. As a reasonable compromise between performance and compute requirements, networks were always trained until a validation set accuracy criterion of 95% was achieved.

### Estimating the energy needed for plasticity

Our approach requires a model of the metabolic energy needed for learning. It is currently not known why the metabolic cost of some forms of plasticity is so high, nor is it known which process is the main consumer. In flies persistent, protein synthesis dependent memory is much more costly than protein synthesis independent forms of memory. One could presume that protein synthesis itself is costly, however it has also been argued that protein synthesis is relatively cheap (Karbowski, 2019). After all, also in the absence of synaptic plasticity there is substantial protein turnover. Examples of other costly processes could be transport of proteins to synapses, actin thread-milling, synaptogenesis. In addition, there might be energy costs that are not related to individual synaptic plasticity, such as replay processes (see Discussion).

We propose a generic model for the metabolic energy *M* of synaptic plasticity under the following assumptions:

1. While it is well known that there are interactions between plasticity of nearby synapses (e.g. Harvey and Svoboda, 2007; Wiegert et al., 2018), we assume that there is no spatial interaction in regards to synaptic costs. It might be that potentiation of two synapses is relatively cheaper than potentiation of a single synapse, e.g. when costly pathways can be shared, as in synaptic tagging and capture (Frey and Morris, 1997; Barrett et al., 2009; Redondo and Morris, 2011). However, instead it is also possible that there is competition for resources (Fonseca et al., 2004; Sajikumar et al., 2014; Jeong et al., 2021), and potentiation of two nearby synapses could be extra costly (Triesch et al., 2018). The interactions when one synapse undergoes potentiation while another one undergoes depression, might even be more complicated as resources might be reused. In summary, both cooperative as well as competitive interactions likely exist but as there is currently no information about their energetics, we need to ignore them. The This implies that the energy is the sum of the individual costs, *M* = ∑*_i_M*^(i)^, where *i* sums over all weights in the network, and *M^(i)^* is the energy required to modify synapse *i*.
2. Similar arguments can be made about the temporal interactions between plasticity. Again, there is extensive literature on temporal interactions in plasticity induction and expression (Petersen et al., 1998; Abraham et al., 2002; Kramár et al., 2012). Energetically, it might be cheaper to potentiate a synapse twice in rapid succession, but it could equally be more costly. We assume that there is no temporal interaction in regards to synaptic costs. This implies that the total metabolic cost is summed over all timesteps *M* = ∑*_t_M* (*t*).
3. Next, we assume that synaptic potentiation and depression both incur the same cost. As we are typically interested in savings in energy between algorithms, this assumption turns out to be minor. Consider a case where, say, synaptic potentiation costs energy but depression does not. As synapses undergo a similar number of potentiation and depression events during training such a variant would halve the cost, however, it does not substantially change the achievable savings, Fig. 6.

Under the assumption of spatial and temporal independence we still require an expression how much modification of a single synapse costs. We propose that this scales as |*δw_i_*|^*α*^, where *α* is a parameter. The total energy sums across all neurons and all training time steps is

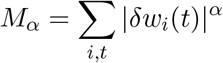

The parameter *α* expresses the proportionality in the weight change. For *α* = 1 the energy (*M*_1_) is linear in the amount of weight change and represents the accumulated weight changes. This is for instance relevant for the energy consumed by protein synthesis where larger changes would require more proteins. Larger values of *α* would lead to a situation where updating a synapse twice would be cheaper than updating once with double the amount. This is not impossible, but seems unlikely.

In the limit *α* → 0, *M_α_* counts the total number of synaptic updates, irrespective of their size. As an example, this would be the case when synaptic tag setting would be costly (Barrett et al., 2009). It is also a reasonable cost function for digital computer architectures, where the cost of writing a memory is independent of its value. While intermediate values of α as well as combinations of terms are possible, we concentrate on *M*_0_ and *M*_1_.

The energy measures are summed across both layers of the network. The number of synapses of both layers is proportional to the number of hidden units. But as dictated by the task, the input-to-hidden layer has 784 synapse per hidden unit, while the hidden-to-output layer has only 10 synapses per hidden unit. In unmodified networks the plasticity cost of the input-to-hidden layer therefore dominates the total energy.

## Results

### Energy of learning

We first explored the energy requirements of training standard large networks. We compared the training of a network with 100, 1000, and 10000 hidden units, Fig.1a. In addition to the number of iteration needed to reach criterion performance (top panel), we measured the two energy variants described in the Methods. The total number of synaptic updates, called *M*_0_, is directly proportional to the learning time. If the learning rate is too low, learning takes too long; too high and learning fails to converge. An intermediate learning rate minimizes the learning time and hence *M*_0_ energy, Fig.1a; middle panel. The optimal learning rate is in good approximation independent of network size.

The cumulative weight change, called *M*_1_ energy, measures the total amount of absolute weight changes of all synapses across training, Fig.1a; bottom panel. This energy measure is smallest in the limit of small learning rates, where it becomes independent of the learning rate. With a small learning rate the path in weight space is more cautious without overshooting associated to larger learning rates. While the energy measures *M*_0_ and *M*_1_ thus have a different optimal learning rate, below we use a learning rate of 0.01 as a compromise, allowing direct comparison.

### Cost of learning in large networks

The MNIST classification networks typically have layers consisting of some hundred units. However, as in biology the number of neurons is enormous (see Discussion), we examine the energy consumed as a function of the network size, Fig.1b. First consider the total number of updates, *M*_0_. Because our setup rules out synaptic updates that are accidentally exactly zero (see Methods), the number of non-zero updates is the total number of synapses in the network multiplied by the number of iterations *T*. Larger networks learn a bit quicker than smaller ones, 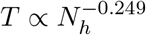, Fig.1c top, where *N_h_* denotes the number of units in the hidden layer. As a result, energy scales sub-linearly with network size 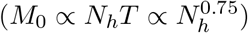.

The accumulated synaptic change energy *M*_1_ also increases for larger networks albeit less steeply, Fig.1b. Despite a fixed learning rate, the size of individual weight updates is smaller in larger networks, Fig.1c, bottom (c.f. Lee et al., 2019). In large networks individual synapses 1) take fewer and smaller steps, and 2) the final individual weights have smaller norm as the larger parameter space of large nets allows solutions closer to the initial weights. We find an approximate square root relation between energy and size, 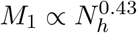, Fig.1b.

### Randomly restricting plasticity to save energy

The above result shows that while large, over-dimensioned networks learn a bit faster, training them uses far more energy – restricting plasticity might save energy. To examine this we first only allow plasticity in a random subset of synapses using a random binary mask. Mask elements were drawn from a Bernoulli distribution. The mask was fixed throughout training. When a different random mask was used at every trial (as in Salvetti et al., 1994), learning simply slowed down and masking did not save energy; also using a different mask for each output class did not save energy.

Using a mask, both energy measures strongly reduced, Fig. 2a, green curves. The optimal mask density was estimated from a counting argument. (We also numerically optimized the fraction of plastic synapses in the input layer; this gave similar results). It is based on the above observations that to achieve the criterion performance one needs at least about 100 hidden units in case that all connections are plastic, Fig. 1b. In other words, there need to be about *μn_in_n_out_* plastic paths between any input and any output, where *n_in_* = 28^2^ is the number of inputs and *n_out_* = 10 the number of output units, and *μ* = 100. We now assume that this is the critical property of the network. According to the hypergeometric distribution, random masks in the input and output layer reduce the mean number of plastic paths to *μ* = *f*_0_*f*_1_*N_h_*, where *f_i_* denotes the fraction of plastic connections in layer *i*.

At the optimal probability, we find that the weight change is approximately the same across network size and layer. Hence both the *M*_0_ and *M*_1_ energy are proportional to *m*(*f*_0_, *f*_1_) = *n_in_N_h_f*_0_+*N_out_N_h_f*_1_, We minimize *m*, subject to *μ* = *f*_0_*f*_1_*N_h_* and 0 ≤ *f_i_* ≤ 1. For a given μ ≥ *N_h_*, the energy is minimal when 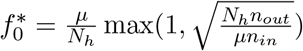 and 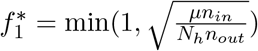, see Fig.2b. Because the number of inputs by far exceeds the number of outputs, it is optimal to keep all hidden-to-output connections plastic (*f*_1_ = 1). Only when 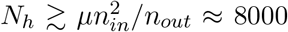, it becomes wasteful to keep all outgoing connections plastic and it is best to reduce the fraction of plastic output synapses as well. The energy scales as

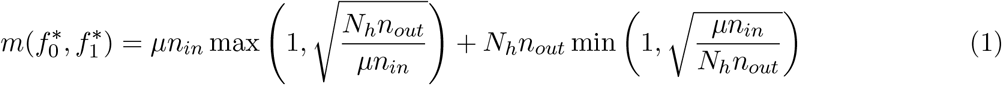

The energy according to this estimate is plotted as the dashed curve in Fig. 2, with the proportionality constant extracted from the value of the energy at *N_h_* = 100. The energy increases in large networks as not every plastic connection in the input-to-hidden layer is met with a plastic connection in the hidden-to-output layer. For large networks the energy scales as 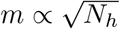.

A similar approach is to mask on a per neuron basis and only allow plasticity in a fixed subset of neurons in the hidden layer. In practice we multiply the vector of back-propagated errors elementwise with a fixed binary mask. The mask density was optimized for each network size and for both energy measures separately. This algorithm saved a bit more energy but in large networks the lack of coordination between plasticity input and output layers, again wastes energy Fig.2a, blue curves. For the *M*_1_ energy the optimal number of plastic units in the hidden layer is around 50…100, irrespective of network size, Fig. 2c. In contrast the *M*_0_ energy was minimal at a lower number of plastic neurons, preferring fewer, but larger synaptic updates.

There is a trade-off between energy saving and learning speed. In unrestricted networks, training is faster is large networks, but unsurprisingly, with all masking variants, training no longer speeds up, Fig.2a inset.

It is interesting to note that for networks with many hidden units (*N_h_* ≳ 5000), it is possible to fix all synapses in the hidden layer, and restrict plasticity to the hidden-to-output synapses, so called extreme learning machines (Schmidt et al., 1992; Huang et al., 2006). However, the plasticity in the hidden-to-output synapses still requires energy and we found that when trained with SGD such a setup did not save more energy than the methods presented here. Moreover, it required a precise tuning of the variance of the input-to-hidden weights.

### Coordination of plasticity: subnets

The above algorithm does save a lot of energy compared to standard backprop but very large networks still require more energy. Inspired by the masking algorithm, we coordinate plasticity between input and output of the neuron: if and only if neuron’s incoming connections are plastic, then so are its outgoing ones. For large networks this is saves energy because such coordination ensures that energy becomes independent of network size, Fig.2a, purple diagram+curves. This has a straightforward explanation: the coordination of incoming and outgoing plasticity results in a plastic sub-network embedded in a larger fixed network. The energy needed for plasticity is the same as in a network with the size of the subnet, hence the presence of the static neurons do not hinder or facilitate the learning. The optimal size of the plastic subnet weakly depends on the energy measure used (about 60 for *M*_0_ and 100 for *M*_1_). As can be inferred from Fig.2a, using, say, 1000 neurons for the subnet does not dramatically increase energy.

### Restricting plasticity to synapses with large updates

In an effort to further decrease energy need, we next modified only synapses with the largest magnitude updates (Golub et al., 2018). Upon presentation of a training sample, the proposed weight updates in the input-to-hidden layer were calculated via standard back-propagation. However, only synapses with the largest magnitude updates (positive or negative) were competitively selected and modified. Synapses in the hidden-to-output layer were always updated. The set of selected synapses was recalculated for every iteration.

The competition was done in two ways: 1) on the level of each neuron so that only fraction of synapses per neuron was plastic, 2) across the whole input-to-hidden layer. It is also possible to select only neurons with the largest activities, or neurons with the largest backpropagation error to have plastic synapses. This also saves energy, but not as much as selecting on weight update per layer. The savings are illustrated in Fig. 3a for a network with 1000 hidden units. Restricting the synaptic updates to those with the largest gradients saves *M*_0_ energy in particular, Fig. 3a.

We wondered if this competitive algorithm in effect works the same as the fixed mask. In other words, does it always select the same synapses to be updated? To examine this we calculated the probability that a given synapses is updated throughout training and extract the inverse Simpson index 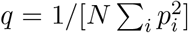 (Hill, 1973), where *p_i_* is the extracted update probability for synapse *i*. When always the same *k* synapses would be updated, *q* = *k/N*. When the updates would be distributed over all synapses, *q* = 1. We find that *q* ≈ 0.6 both at the start and end of training, which is much larger than *k/N* ≈ 0.01. In contrast to the fixed masks used above, plasticity keeps switching between synapses under this algorithm.

Fig. 3b examines the savings across network size, showing large *M*_0_ but little *M*_1_ savings. The reason for this can be seen in the diagram Fig. 3c. The *M*_1_ energy, i.e. L1 path length, is independent of the weight trajectory when it does not backtrack (top row). Only when the weight paths backtrack can *M*_1_ be saved (middle row), but even this is not guaranteed (bottom row).

Next, we further sought to decrease energy need by selecting only the synapses with the largest updates in the subnet. This is the same algorithm as above but now restricted to the subnet. This further decreased the energy needs and again made them independent of network size, Fig.3 (purple curve).

In summary, the most energy efficient plasticity occurs when 1) on a given sample only the largest updates are implemented, and 2) these synapses form a plastic pathways in the network. The competitive selecting via the maximum has another advantage over a fixed mask. While the fixed mask limits the capacity of the network, the maximum selection does not. To show this we used a small network (100 hidden units) and trained for 30 epochs, at which time accuracy had saturated. When using a fixed mask, the restricted network’s performance drops with mask size, Fig.3d. However using either competitive selection mechanism, the maximum accuracy dropped much less.

### Combining mask with synaptic caching

It might appear that we have exhausted all ways to save plasticity energy, but additional savings can be obtained by reducing the inefficiency across trials. During learning the synaptic weights follow a random walk, often partly undoing the update from the previous trial. This is inefficient.

An additional, different way to save energy consumed by plasticity exploits that not all forms of plasticity are equally costly. Transient changes in synaptic strength (early phase LTP in mammals, ARM in fruit-flies) are metabolically cheap, while in contrast permanent changes (late-phase LTP in mammals, LTM in fruit-flies) are costly (Mery and Kawecki, 2005; Plaçais and Preat, 2013; Potter et al., 2010). It is possible to save metabolic cost by storing updates initially in transient forms of plasticity and only occasionally consolidate the accumulated changes using persistent plasticity. We have termed this saving algorithm *synaptic caching* (Li and Van Rossum, 2020). It is somewhat similar to batching algorithms in machine learning.

By distributing plasticity over transient and persistent forms, synaptic caching can save energy. The amount of savings depends on how rapid transient plasticity decays relatively to the sample presentations, and its possible metabolic costs. The largest saving can be achieved when transient plasticity comes at no cost and decays so slowly that all consolidation can be postponed to a single consolidation step at the end of learning.

To explore whether the algorithms introduced in this study are compatible with synaptic caching we implemented both transient and persistent forms of plasticity in the network coded in *w*^trans^ and *w*^pers^. The total connection strength between neurons was *w*^trans^ + *w*^pers^. Updates of the weights were first stored in the transient component. Only when the absolute value of the transient component reached a threshold value, the transient weight was added to the persistent weight and the transient component was reset to zero. The decay rate of the transient weights was 10^-3^ per sample presentation; the cost of transient plasticity was set to 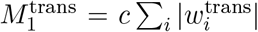, with constant *c* = 0.01 (see Li and Van Rossum, 2020 for motivation and further parameter exploration).

Fig. 4 shows the energy measures as a function of the consolidation threshold. Synaptic caching saves a substantial amount of energy. As the transient plasticity does not contribute to the *M*_0_ energy measure, it now just counts the number of synaptic consolidation events. It is lowest at a high consolidation threshold, at even higher thresholds (≳ 0.06) the learning no longer converges. By thus limiting and postponing consolidation the *M*_0_ energy with synaptic caching is virtually independent of network size.

However, the *M*_1_ energy still increases with network size, with a similar dependence on network size as the standard network, Fig. 4b. It will still lead to high costs in large networks. Coordinating both transient and persistent plasticity, that is restricting plasticity to subnets, again eliminates this increase. We also tried a variant in which transient plasticity was distributed over the whole network and only consolidation was coordinated, however this did not save energy.

In sum, combining synaptic caching with the above savings strategy saves the most energy. For the *M*_0_ energy synaptic caching even completely obviates the need for additional saving strategies in large networks. We emphasize however that synaptic caching does rely on additional synaptic complexity, namely two synaptic plasticity components and a consolidation mechanism. Moreover, if the updates of the transient component would also incur *M*_0_ energy, additional saving strategies might be possible.

## Discussion

Experiments have shown that already simple associative learning is metabolically costly. For neural networks that learn more complicated tasks, energy costs can become very high, in particular when networks are large. For instance, macaque V1 has some 150 million neurons and some 300×10^9^ synapses (Vanni et al., 2020). Extrapolating Fig.1, if plasticity were distributed over all these synapses, the number of synaptic updates would be some 10^5^ times larger than required. The *M*_1_ energy, which increases less steeply with network size, would still be some 700 times larger, Fig.5. Thus large networks are powerful, but without restrictions the metabolic cost of plasticity could be very large.

We have introduced two approaches to reduce costs. First, restrict plasticity to a subset of synapses.

This is most efficient when the plasticity on input and output side of a neuron are coordinated, so that when inputs of a neuron are modified then so are its outputs. Such effects have indeed been observed, although the precise rules appear complex (Fitzsimonds et al., 1997; Tao et al., 2000; Zhang et al., 2021).

Second, express plasticity only in synapses with large updates. This is also consistent with neuroscience data, where there is typically a threshold for plasticity induction. Our study would suggests the presence of a competitive algorithm in which only a certain fraction of synapses is modified. Bio-physically, the competition on the neuron level could naturally follow from resource constraints.

These strategies can be combined with our earlier work on synaptic caching. Finally it also possible to skip over uninformative samples during learning, leading to even further saving (Pache and van Rossum, 2023).

Future studies could include how the saving algorithms should be automatically adapted dependent on task difficulty or network architecture. Another interesting avenue is to find the most energy efficient synaptic modification during transfer learning (cf. von Oswald et al., 2021).

Given the lack of biological data and the uncertain nature of the main energy consumer, our proposed energy model is currently coarse. In particular the independence assumption is unlikely to be fully correct; yet even the sign of the interactions (cooperative vs competitive) is unknown. We assumed that the energy is extensive in the number of synapses. It might be that there is a large component independent of synapse number, for example energy needed for replay processes. In that case, the question is at what number of synapses the energy becomes approximately extensive. As more experimental data becomes available, the energy model can be refined and the efficiency of the proposed algorithms can be re-examined.

Finally, we did not consider the energy required for neural signaling, such as synaptic transmission and spike generation. Ultimately, learning rules should aim to also reduce those costs.

## Acknowledgments

It is a pleasure to thank Mikhail Belkin, Simon Laughlin, Thomas Oertner, Aaron Pache, Joao Sacramento, Long Tran-Thanh, and Silviu Ungureanu for discussion. This research was supported by a grant from NVIDIA and utilized an NVIDIA RTX A6000.

## Appendix

